# Proprioceptive and visual feedback responses in macaques exploit goal redundancy

**DOI:** 10.1101/2022.02.11.480080

**Authors:** Kevin P. Cross, Hui Guang, Stephen H. Scott

## Abstract

A common problem in motor control concerns how to generate patterns of muscle activity when there are redundant solutions to attain a behavioural goal. Optimal feedback control is a theory that has guided many behavioural studies exploring how the motor system incorporates task redundancy. This theory predicts that kinematic errors that deviate the limb should not be corrected if one can still attain the behavioural goal. Several studies in humans demonstrate that the motor system can flexibly integrate visual and proprioceptive feedback of the limb with goal redundancy within 90ms and 70ms, respectively. Here we show monkeys (*Macaca mulatta*) demonstrate similar abilities to exploit goal redundancy. We trained four male monkeys to reach for a goal that was either a narrow square or a wide, spatially redundant rectangle. Monkeys exhibited greater trial-by-trial variability when reaching to the wide goal consistent with exploiting goal redundancy. On random trials we jumped the visual feedback of the hand and found monkeys corrected for the jump when reaching to the narrow goal and largely ignored the jump when reaching for the wide goal. In a separate set of experiments, we applied mechanical loads to the monkey’s arm and found similar corrective responses based on goal shape. Muscle activity reflecting these different corrective responses were detected for the visual and mechanical perturbations starting at ∼90 and ∼70ms, respectively. Thus, rapid motor responses in macaques can exploit goal redundancy similar to humans, creating a paradigm to study the neural basis of goal-directed motor action and motor redundancy.

## Introduction

A common problem in motor control concerns how to select muscle commands when there are many different ways to attain a behavioural goal (Bernstein, 1967; Flash and Hogan, 1985; Sporns and Edelman, 1993; Scholz and Schöner, 1999; Scholz et al., 2000; Latash, 2012). For example, successfully reaching for an object can involve many different trajectories of the limb to the goal, and thus, many different patterns of muscle activity. Optimal feedback control (OFC) provides a framework for how to select motor commands among a family of redundant solutions (Todorov and Jordan, 2002; Scott, 2004). OFC selects motor commands to optimize a cost function that balances successfully completing the behavioural goal with the cost of movement (e.g. energy, noise). Importantly, these controllers abide by the minimum intervention principle where kinematic errors that arise during movement are only corrected if they interfere with the behavioural goal. Or alternatively stated, errors that deviate the plant along redundant trajectories should not be corrected. As a result, variability accumulates along redundant task dimensions.

Several studies demonstrate that the motor system exploits task redundancies similar to OFC controllers (Diedrichsen, 2007; Dimitriou et al., 2012; Cluff and Scott, 2015; Weiler et al., 2015, 2016). A common approach is to have participants reach to a spatially redundant goal such as a wide rectangular bar (Knill et al., 2011; Nashed et al., 2012; de Brouwer et al., 2017; Keyser et al., 2017, 2019). Participants exhibit greater trial-to-trial variability in their reach endpoints when reaching for a wide as compared to a narrow goal. The increased variability also exhibits structure as variability primarily accumulates along the wide (redundant) axis of the goal. Furthermore, displacements to the visual feedback of the hand (cursor jump) are fully corrected by participants when reaching for a narrow goal and are corrected less when reaching for the wide goal if the displacement is along the redundant axis (Knill et al., 2011; de Brouwer et al., 2017; Cross et al., 2019). Differences in these corrective responses arise in muscle activity ∼90ms after the jump (Franklin and Wolpert, 2008; Cross et al., 2019). Similar corrective responses occur when mechanical loads are applied to the limb with differences between corrective responses starting ∼70ms after the load (Nashed et al., 2012; Lowrey et al., 2017; Keyser et al., 2019). However, despite the prevalence of OFC as a theory of motor control, we know little about the neural circuits used to generate OFC-like behaviours.

One challenge with investigating neural circuits underlying rapid motor responses is that behavioural tasks must be translated into animal models that allow for invasive neural recordings such as rhesus monkeys or rodents. To perform most behavioural tasks the behaviour of the animal must be shaped using reward over the course of tens of thousands of trials. This excessive training along with behavioural shaping results in behaviour that is highly reproducible on a trial-by-trial basis which provides advantages when analyzing noisy neural activity. However, it is unclear if this highly reproducible behaviour comes at the cost of behavioural flexibility. For example, Bizzi and colleagues trained monkeys to reach to a goal and applied an assistive mechanical load that pushed the monkey’s limb towards the goal (Bizzi et al., 1982, 1984). Monkeys corrected by actively resisting the load in what appeared to be an attempt to return the limb to the original trajectory to the goal. In contrast, OFC models and humans performing a similar task do not return to the original trajectory and instead allow the load to push the hand towards the goal thereby exploiting the fact that there are redundant trajectories (Cluff and Scott, 2015). Thus, humans can exploit task redundancies, however it is unclear whether monkeys are capable of exploiting redundancies in an experimental setting due to overtraining and behavioural shaping.

Here, we investigated whether monkeys could learn to exploit the spatial redundancy of a goal during reaching. We found monkeys exhibited greater variability in their reach endpoints on unperturbed trials. Further, monkeys corrected less for visual and mechanical perturbations of the limb when reaching for the spatially redundant target, consistent with exploiting goal redundancy. Muscle recordings indicated that feedback responses to visual and mechanical perturbations reflected goal redundancy within <100ms.

## Methods

Four male monkeys (*Macaca mulatta*, 10-20kgs) were trained to sit in a primate chair and place their upper arm into a robotic exoskeleton (Scott, 1999, Kinarm, Kingston, Canada). The robot constrained the monkey’s arm to move in a two-dimensional plane and included a virtual reality system that could display virtual targets and visual feedback of the limb. Experiments were approved by the Queen’s University Research Ethics Board and Animal Care Committee.

### Behavioural task for visual perturbations

Monkeys were trained to make goal-directed reaches to targets in the virtual environment. Visual feedback of the hand was provided by a white cursor (radius=0.8cm) aligned with the monkey’s index fingertip. At the start of each trial, a start target (square, side length=1.2cm) appeared and the monkey was required to reach and hold their hand at the target for 750-1500ms. Next, a goal target appeared that was located 8cm lateral, and 4.4cm in front of the start goal (total reach distance 9.2cm, Figure 1A). To reach the goal from the starting position monkeys had to primarily extend their elbow. When the monkey’s hand left the start target, they had 900ms to reach and stabilize their hand within the goal for 500ms. The goal could either be a narrow rectangle (length 2cm, width 2.2cm) or a wide rectangle with its long axis oriented perpendicular to the reach axis (straight path between start and middle of goal target; wide goal dimensions length 12cm width 2.2cm). On random trials, the cursor jumped orthogonal to the reach axis (orthogonal reach axis) once the hand was 2cm from the start target and on these trials monkeys were given an additional 500ms to reach to the goal. For monkeys M and A, the cursor jumped ±4cm and for monkeys T and C the cursor jumped ±3.5cm. Within a block of trials there were 6 no-jump reaches (3 for both goal shapes), and 4 cursor-jump trials (2 directions x 2 goal shapes). Monkeys completed 10-25 blocks in a given recording session.

**Figure 1.**
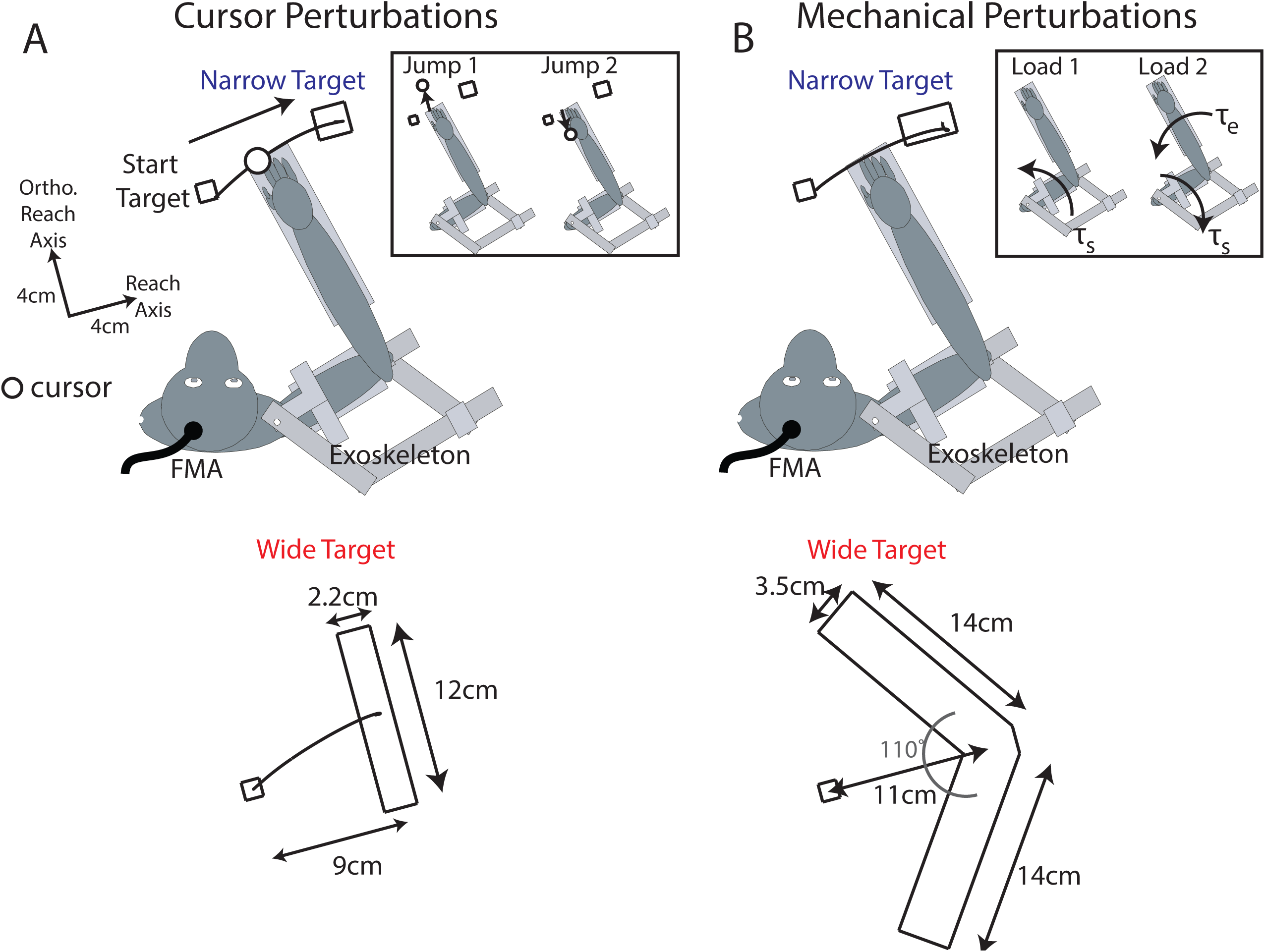
Experimental set-up. A) Monkeys placed their arm inside a robotic exoskeleton and were trained to reach from a starting position (Start Target) to either a narrow target (top) or wide target (bottom). Inset: on random trials, the visual feedback of the hand (cursor) jumped orthogonally by 3-4cm (see Methods). B) Same as A) showing the configuration of the targets during the mechanical perturbation experiment. Inset: on random trials, mechanical loads were applied to the limb that displaced the limb orthogonally.

### Behavioural task for mechanical perturbations

The task was similar to the cursor perturbation task, but there were several changes to the shape and size of the goal targets. First, the size of the redundant axis of the wide goal was 28cm long (Figure 1B) and it was shaped like an arrowhead composed of two rectangles overlapping at the edges and at an angle of 110° with respect to each other. We chose this configuration as it brought the edges of the goal closer in proximity to the monkey’s arm, thus making it easier to reach on perturbation trials. On random trials mechanical loads were applied to the joints to displace the limb approximately orthogonal to the reach axis (Monkey M|A|T|C pushes limb away from body: shoulder 0.5|0.45|0.4|0.3 Nm elbow 0|0|0|0Nm; pushes limb towards the body: shoulder -0.5|-0.45|-0.4|-0.3Nm elbow 0.15|0.1|0.07|0.14Nm; + flexion loads, - extension loads). Due to the influence of limb mechanics (i.e. limb inertia, limb geometry), loads were adjusted for each monkey to generate roughly similar hand deviations. Cursor feedback was temporarily removed for 200ms when the load was applied. We also increased the number of unperturbed trials to reduce anticipatory corrections to the mechanical loads. Within a block of trials there were 8 no-load reaches (4 for both goal shapes), and 4 load trials (2 directions x 2 goal shapes). Monkeys completed 10-15 blocks in a given recording session.

### Estimating visual onsets

On cursor-jump trials, there is approximately a 20-49ms delay between when the command was sent to jump the cursor and when it actually updated the screen. We estimated this delay on a trial-by-trial basis by using photodiodes placed at the side of the screen and flashing white squares coincident to the photodiode locations when the cursor jumped.

### Muscle recordings

We implanted Monkey M with a 32-channel chronic EMG system (Link-32, Ripple Neuro, Salt Lake City UT). The system had 8 leads that were inserted into the muscle belly and were attached to a processor. Each lead had 4 separate contacts for recording intramuscular activity (impedance 20 kOhms). The processor was implanted under the skin and located near the midline of the back at the mid-thoracic level. Muscles implanted were brachioradialis, brachialis, the lateral and long heads of the triceps, biceps (long head), pectoralis major, and anterior and posterior deltoids. An external receiver was secured to the skin over the processor using magnets in the receiver and processor. The external receiver was capable of powering the internal processor and EMG signals were transmitted through the skin from the processor to the receiver by photodiodes. The signals were then relayed to the Grapevine Neural Interface Processor (Ripple Neuro, Salt Lake City, UT), bandpass filtered (15-375Hz) and recorded at 2kHz.

In Monkey C we recorded muscle activity using surface EMG electrodes (Delsys, Natwick MA, USA). We recorded activity from brachalis, biceps, and the lateral and long head of the triceps. Activity was sampled at 1kHz, bandpass filtered at 20-200Hz using a 6^th^ order Butterworth filter and rectified.

### Kinematic recordings

The shoulder and elbow angles, angular velocities and angular accelerations were recorded by either a 128-Channel Neural Signal Processor (Blackrock Microsystems, Salt Lake City, UT) at 1kHz, or by the Grapevine Neural Interface Processor at 30 kHz.

## Data and statistical analyses

### Kinematic analysis

The endpoint position of the reach was calculated by finding the hand position at the end of the trial (timepoint after 500ms hold period). This hand position was then projected onto the redundant axis of the wide goal by finding the location on the redundant axis that had the shortest distance to the hand position. The endpoint position was also made relative to the center of the relative target as there were small positional offsets between the narrow and wide targets.

We estimated the timing of the kinematic corrections using the hand velocity along the orthogonal reach axes, which was defined as the velocity component perpendicular to the straight path connecting the start and middle of the goal (reach axis, Figure 1). The hand velocity was aligned to the onset of the perturbation (cursor jump or mechanical load) or the equivalent time point on no-perturbation trials (faux jump or faux load). The average velocity on no-perturbation trials was subtracted from the velocity on perturbation trials resulting in the change (Δ) in hand velocity. Timing of when the hand velocity differentiated based on goal shape was determined using receiver-operator characteristics (ROC) (Corneil et al., 2004; Gu et al., 2016; Pruszynski et al., 2016; Cross et al., 2019). At each time point we generated an ROC curve between the hand velocities for the narrow and wide goals. The area under the ROC curve reflects how discriminable the trials for the narrow and wide goals are and can range from 0.5, indicating chance discrimination, to 0 and 1 indicating perfect discrimination. We found the first time point that had an area >0.8/<0.2 and that was maintained above this threshold for 10 consecutive time points (10ms). We then traced backwards in time to the first time point that fell below/above 0.6/0.4 (knee).

### EMG analysis

For indwelling muscle activity recorded from Monkey M we down sampled activity to 1kHz. For each lead, we computed differential signals from the two most proximal contacts and two most distal contacts relative to the processor. For muscle activity recorded using surface electrodes from Monkey C, we only included recordings where perturbation-related activity was detected.

The differential signals were rectified and low-passed filtered with a 6^th^ order zero-phase lag Butterworth filter at 50Hz. Muscle activities were aligned to the onset of the perturbation. Muscle activities were then trial-averaged and the activities on no-perturbation trials were subtracted from perturbation trials to yield the change in activity caused by the perturbation. Muscle activities were normalized by the mean perturbation-related activity from 0-300ms after perturbation onset.

### Analysis of no-perturbation trials

We compared magnitudes of muscle activity between reaches for the narrow and wide goals on no-perturbation trials. Activities were averaged in the epoch starting 200ms before the faux-jump/faux-load onset until 200ms after the onset (movement epoch). A two-sample t-test identified muscles that were significantly different between the narrow and wide goals (p<0.01). We also examined how temporally correlated muscle activities were between the narrow and wide goals during the movement epoch using Pearson’s correlation coefficient. We compared the observed distribution across muscles with a shuffled distribution where correlations were computed between randomly selected muscles.

### Preferred direction and perturbation-sensitive criteria

A muscle’s preferred perturbation direction was calculated by averaging the perturbation-related activities over the first 300ms after the perturbation onset for reaches to the narrow goal. The direction with the largest activity over this epoch was defined as the preferred direction. The same preferred direction was used for both the narrow and wide goals.

Perturbation-sensitive muscle samples were identified using a two-sample t-test comparing the activity on unperturbed trials with activity on perturbation trials in the muscle’s preferred direction in the epoch of 0-300ms after the perturbation onset. This was applied twice for each muscle, one test for each target shape. Muscles were classified as “perturbation sensitive” if p<0.05 (Bonferroni correction factor 2).

### Epoch analysis

We compared how perturbation-related activities differed between goal shapes over time. For the mechanical perturbations we divided each muscle’s activity into epochs of 20-50ms, 50-75ms, 75-100ms and 120-180ms based on previous work (Lee and Tatton, 1975; Crago et al., 1976; Bonnet, 1983; Omrani et al., 2014; Pruszynski et al., 2014). For the cursor perturbations we considered the same epochs but shifted 50ms forward in time (70-100ms, 100-125ms, 125-150ms and 170-230ms) to account for the fact that muscle activities respond to visual feedback ∼50ms slower than for proprioceptive feedback (Franklin and Wolpert, 2008; Pruszynski et al., 2010, 2016; Dimitriou et al., 2013; Cross et al., 2019). For each muscle, we applied a two-way ANOVA with time (4 levels) and goal shape (2 levels) as factors. Muscles had significant goal-shape modulation if there was a significant main effect for goal shape or if there was an interaction effect (p<0.05 Bonferroni correction 2). Significant interaction effects were decomposed using post-hoc t-tests by comparing across goal shape within each time epoch (p<0.05 Bonferroni correction 4).

### Population signal

We calculated the population signal by averaging the activities for all perturbation-sensitive muscle samples in their preferred directions. A difference signal was calculated by subtracting the activities for the wide goal from the narrow goal for all perturbation-sensitive muscle samples followed by averaging across samples. The onset for the population signals were estimated by calculating the mean and standard deviation (SD) of the baseline activity (300ms before perturbation onset) and finding the first time point that exceeded the mean by 3SD for 20 consecutive time points (Omrani et al., 2016).

## Results

Our goal was to develop a reaching task to examine if monkeys could exploit the spatial redundancy of a goal. The original tasks performed by humans involved a multi-joint reach directly in front of the shoulder joint (Nashed et al., 2012; Cross et al., 2019). However, our initial attempt to translate this task into the monkey was not successful as the monkey exhibited substantial bias towards reaching one end of the bar (note: humans can also show a similar bias, albeit smaller, see Keyser et al., 2017). Instead, we focused on reaches that primarily required an elbow extension movement, which produced more consistent behaviour (Figure 1). Below, we first describe the experiments that involved cursor perturbations followed by the experiments that involved mechanical perturbations.

## Experiment 1: Goal redundancy and feedback responses to cursor jumps

We trained four monkeys to reach to a goal that could be either a narrow square (Figure 1A top) or a wide rectangle (Figure 1A bottom; trials interleaved). For monkeys M|A|T|C, we recorded 6|7|14|12 behavioural sessions of the animals performing the task on separate days which were spread across 1-2 years for each monkey and monkeys were able to perform the task with high efficiency (success rates monkeys M|A|T|C: narrow targets=95|98|89|97%, wide targets=96|100|92|98%).

Figure 2A shows the hand paths for monkey M to the narrow (left) and wide (right) goals from one recording session. There was greater trial-by-trial variability in the hand position during the reach to the wide goal, whereas variability was considerably smaller during reaches to the narrow goal. This resulted in the reach endpoints exhibiting greater dispersion along the redundant axis of the goal for reaches to the wide goal compared with reaches to the narrow goal (Figure 2A-C). Across recording sessions, there was no significant difference in mean endpoint position for monkeys M and C, whereas there was a 0.7cm and 1.3cm difference between endpoint positions for the narrow and wide goals for monkeys A (t(6)=3.8, p=0.01; Figure 2D) and T (t(13)=9.5, p<0.001), respectively. In contrast, the standard deviation of the reach endpoints were 4.2, 1.5, 2.0 and 1.9 times greater for the wide goal than the narrow goal for monkeys M, A, T, and C, respectively (Figure 2E; paired t-test Monkey M|A|T|C t(5)=8.3|t(6)=4.6| t(13)=6.8| t(11)=10.9, p<0.001|p=0.004| p<0.001| p<0.001).

**Figure 2.**
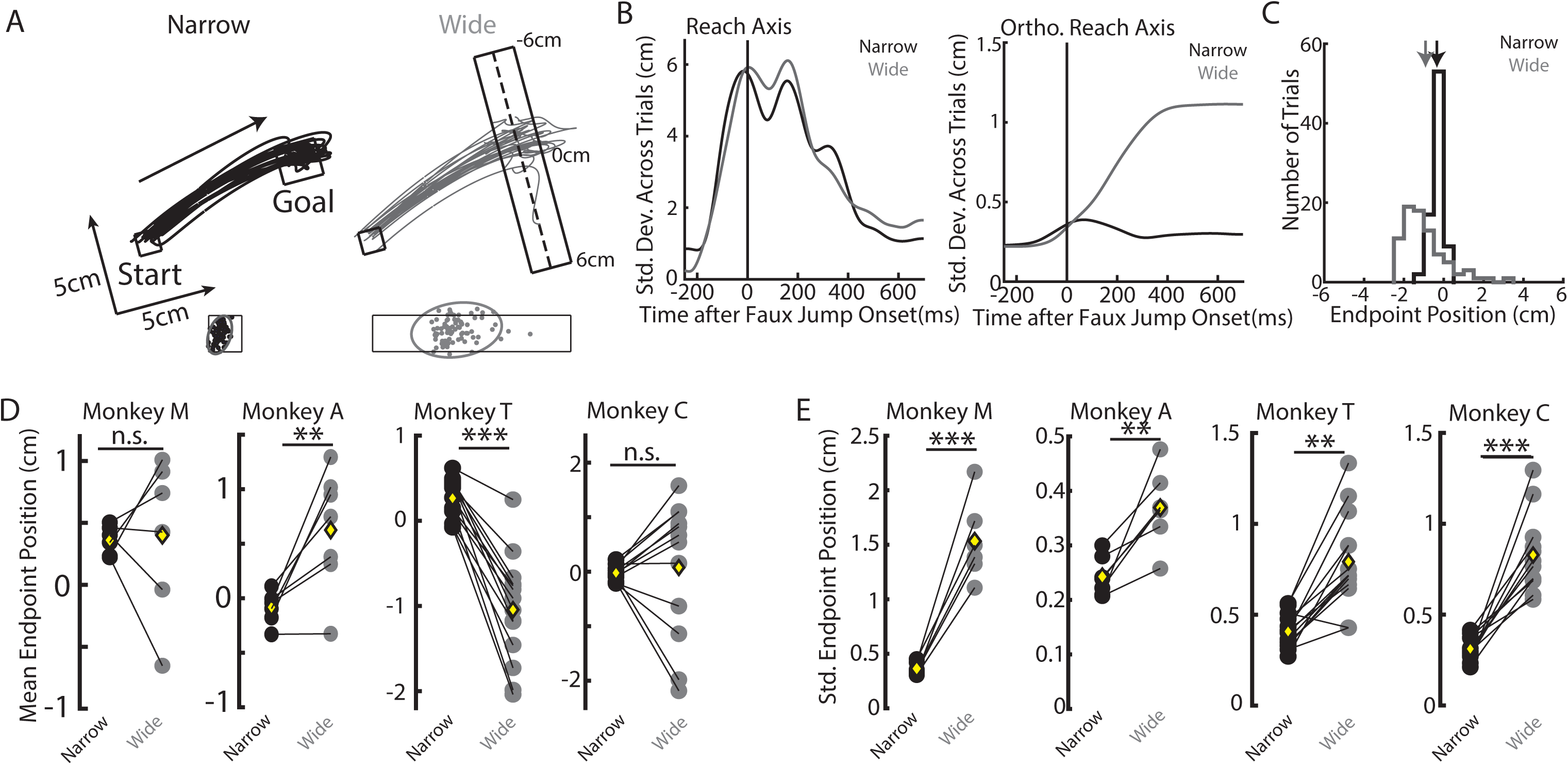
Example kinematics on unperturbed trials for the cursor-jump variant of task. A) Top: Hand paths from one session for Monkey M to the narrow (left) and wide (right) goals. Arrow denotes the direction of reach. Dashed line on the wide goal denotes the redundant axis. Bottom: Reach endpoints and the 95% confidence ellipse. B) Standard deviations of the hand position across trials for the narrow and wide goal reaches. C) Reach endpoint histograms for reaches to the narrow and wide goals from the same session as A). Zero denotes the middle of the redundant axis (A, middle of dashed line). Arrows denote the means of the distributions. D) The mean endpoint position for the narrow and wide goals across all recording sessions. Yellow diamonds denote the means across sessions. E) Same as D) for standard deviation of the endpoint position. Hand paths on cursor-jump trials from the same session as A). ** p<0.01, ***p<0.001.

Next, we examined the unperturbed hand velocities for reaches to the narrow and wide goals. Figures 3A and B (inset) show the unperturbed hand velocities during the same recording session along the reach and the orthogonal reach (ortho. reach) axes, respectively. There was a small increase in the peak hand velocity for reaches to the wide goal as compared to reaches to the narrow goal. Across recording sessions, we found a small increase in hand speed for the wide goal for monkey M (narrow=0.48m/s, wide=0.51m/s, paired t-test t(5)=3.6, p=0.02), a small decrease for the wide goal for monkey T (narrow=0.35m/s, wide=0.32m/s, t(13)=2.5, p=0.02), and no effect of goal shape on hand speed for monkeys A and C (Monkey A|C: narrow=0.52|0.49m/s and wide=0.52|0.5m/s, t(6)=2.0|t(11)=0.2, p=0.1|p=0.8).

**Figure 3.**
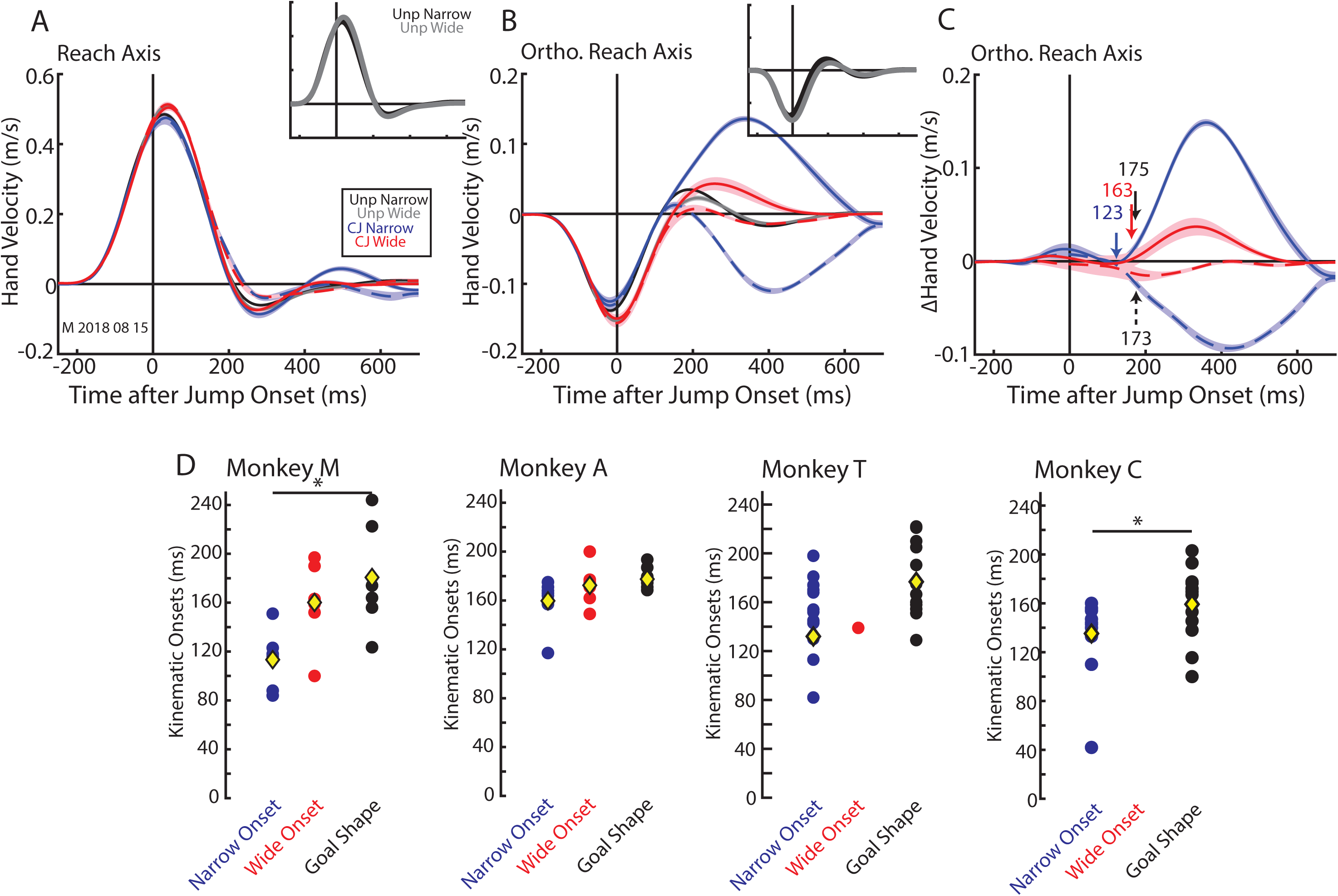
Example hand velocity profiles for the cursor-jump variant of task. A) The hand velocity along the reach axis (see Figure 1) for the narrow and wide goals from the same recording session as Figure 2A). Velocity was aligned to the jump onset. Inset shows the unperturbed reaches to the narrow and wide goals. Unp Narrow and Unp Wide: Unperturbed reaches to the narrow and wide goals, respectively. CJ Narrow and CJ Wide: Cursor jump trials for reaches to the narrow and wide goals, respectively. Solid and dashed lines denote cursor jumps away from the body and towards the body, respectively. B) Same as A) for the hand velocity along the orthogonal reach axis. C) The change in the hand velocity on cursor-jump trials for the narrow and wide goals. Same recording session as A). Blue and red arrows denote the earliest corrective onset for the narrow and wide goals, respectively. Black arrows denote when corrective movements differentiate based on goal shape. D) The earliest corrective onsets for the narrow (blue) and wide goals (red) along with the onsets for when corrections differentiated based on goal shape (black) across sessions. Yellow diamonds are the means across sessions. * p<0.05.

Next, we examined how goal redundancy affected corrective responses for unexpected cursor jumps. Figure 4A shows monkey M’s hand paths on cursor-jump trials for the narrow and wide goals. There is a clear correction for the cursor jump when reaching for the narrow goal and little if any correction when reaching for the wide goal. There was also greater trial-by-trial variability in the hand position during the reach to the wide goal (Figure 4B), indicating that the monkey did not simply learn to reach for a particular location on the goal during cursor-jump trials. We calculated the differences between reach endpoints on cursor-jump trials and the mean of the unperturbed reach endpoint (change in reach endpoint, Figure 4C). For the narrow goal, the change in reach endpoints for either cursor jump direction (blue solid line: jumps away from body; blue dashed line: jumps towards body) were narrow distributions centered near zero (mean: solid 0.26cm, dashed -0.55cm) indicating that monkeys ended their reach at almost the same location as on unperturbed trials. In contrast, for the wide goal the change in reach endpoints generated wide distributions that were centered ∼3cm from the zero-mark (red solid 2.9cm, red dashed -3.6cm) indicating that the monkey largely ignored the cursor jump. Across sessions and in both jump directions, there was a greater change in endpoint position for the wide goal than the narrow goal (Figure 4D, F; paired t-tests: jumps away from body | towards body monkey M t(5)=7.6| t(5)=11, monkey A t(6)=49| t(6)=48.9, monkey T t(13)=33|t(13)=19, monkey C t(11)=24, |t(11)=23.8, p<0.001 for all comparisons). Furthermore, the standard deviation in reach endpoints was also significantly greater for reaches to the wide goal for monkey M (Figure 4E, G; jumps away from body|towards body: paired t-test: t(5)=7.9| t(5)=8.4, p<0.001| p<0.001), monkey T (t(13)=6.4|t(13)=5.6, p<0.001|p<0.001) and monkey C (t(11)=8.4|t(11)=7.1, p<0.001|p<0.001). For monkey A, a significant increase in the standard deviation in reach endpoints was found for jumps towards the body (t(6)=8.6 p<0.001) but not for jumps away from the body (t(6)=1.6 p=0.2). Thus, reach endpoints on cursor-jump trials were more variable and biased towards the edges of the goal during wide goal reaches as compared to narrow goal reaches.

**Figure 4.**
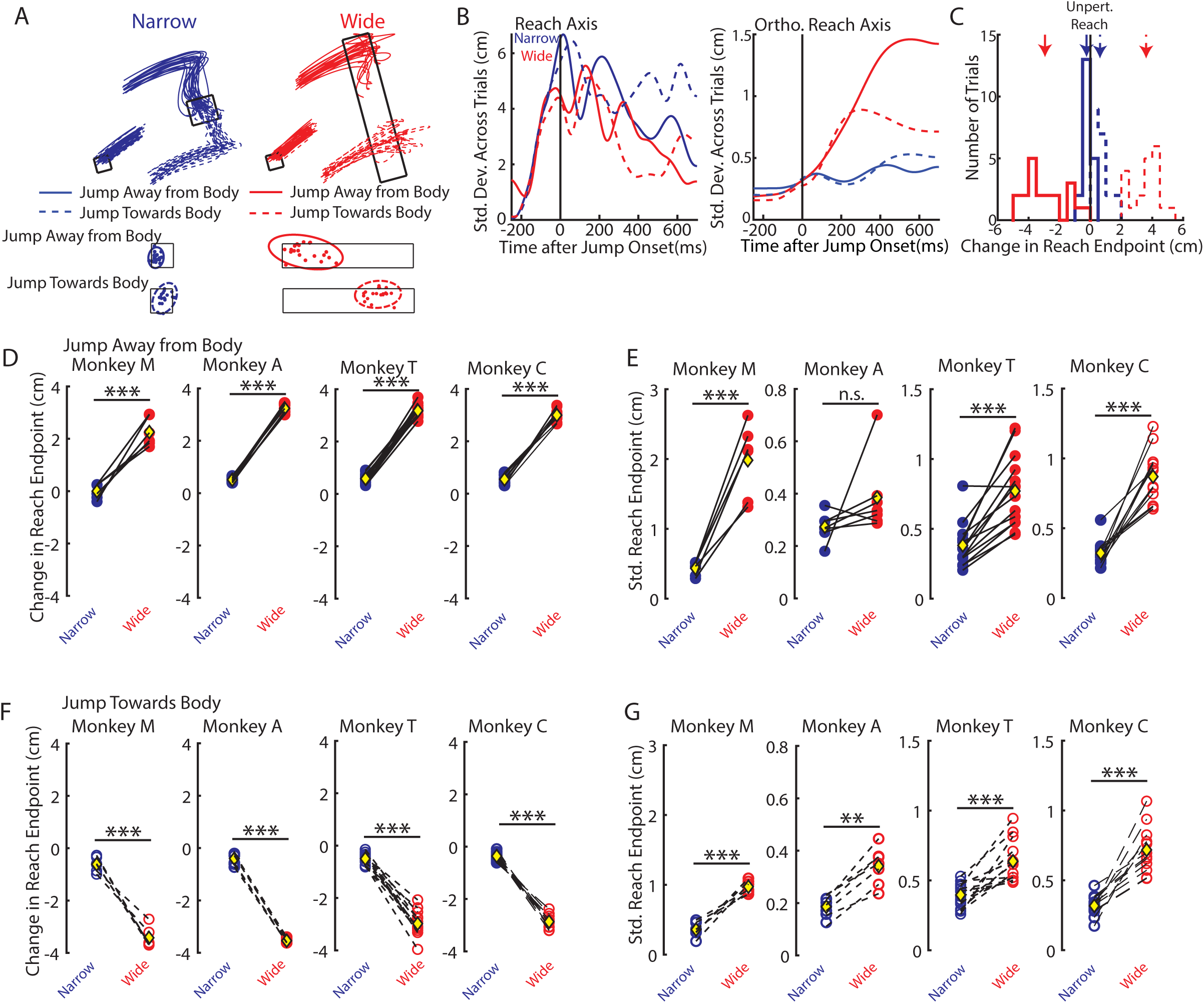
Example kinematics on cursor-jump trials. A) Top: Hand paths from Monkey M for cursor-jump trials. Same session as Figure 2A. Bottom: Reach endpoints and the 95% confidence ellipse. Solid and dashed lines denote cursor jumps away from the body and towards the body, respectively. B) Standard deviation of the hand position across trials for the narrow and wide goal reaches. C) The change in reach endpoint histograms (change relative to unperturbed reach trials). Zero denotes the mean of the reach endpoints on unperturbed trials (Figure 2C arrows). Arrows denote the means of the distributions. D) The mean change in reach endpoint for jumps away from the body across all recording sessions. E) Same as D) for the standard deviation in reach endpoints. F-G) Same as D-E) for jumps towards the body. ** p<0.01, ***p<0.001.

For cursor-jump trials, kinematic changes were primarily restricted to the orthogonal reach axis, which coincided with the direction that the cursor jumped (Figure 3A, B). Figure 3C shows the hand velocity after subtracting the unperturbed hand velocity (Δ ortho reach velocity). For the narrow and wide goals, the monkey initiated a correction for the cursor jump within 123ms (blue arrow, narrow onset) and 163ms (red arrow, wide onset) of the jump onset, respectively. Differences between corrective responses for the narrow and wide goals began to differentiate ∼174ms after the jump (solid and dashed black arrows; goal shape onset). Across recording sessions, monkey M|A initiated a correction within 113|160ms and 160|173ms of the cursor jumps to the narrow and wide goals, respectively. Monkeys C|T initiated a correction for the narrow goal 132|135ms after jump onset but did not reliably detect a correction for the wide goal. Hand velocity differentiated based on goal shape starting at 181|178|176|159ms (average across both directions, Figure 3D) for monkey M|A|T|C. A one-way repeated-measures ANOVA with onset type as a factor (3 levels, narrow and wide goals, and goal shape) was significant for monkey M (F(2,10)=5.0, p=0.03). Post hoc paired t-tests revealed a significant difference between the narrow onset and the goal-shape onset (t(5)=4.1, p=0.02, Bonferroni correction factor of 3). The ANOVA was not significant for monkey A (F(2,12)=2.3, p=0.14). Since the wide goal onsets were largely absent for monkeys T and C, we instead used paired t-tests between the narrow and goal shape onsets and found onsets were not significant (monkey T|C t(13)=2.0| t(11)=2.0, p=0.06|p=0.07).

From monkey M, we implanted an indwelling chronic EMG system that recorded the activities of eight muscles that spanned the shoulder and elbow joints. The system allowed us to sample activity from each muscle twice and allowed us to record the same muscle across multiple recording sessions (see Methods). We recorded muscle activities across two sessions and pooled trials. Since we were largely interested in perturbation-related activity, we restricted our analysis to 14 of 16 samples that responded to the cursor jump (see perturbation-sensitive criteria in Methods). For monkey C, we recorded muscle activity using surface electrodes over the course of four sessions. We only analyzed recording days where perturbation-related activity was detected in a given muscle (number of good sessions: TLat: 3 Br: 4 Bi: 1 TLong: 3), due to the poorer signal-to-noise ratio of surface EMG than indwelling EMG and due to the variability inherent in day-to-day electrode placements.

Figure 5A shows the average activity on unperturbed trials for the pectoralis major (PM shoulder flexor) muscle aligned to the faux-jump onset from monkey M. The temporal structure of the muscle activity was comparable between goal shapes, however, starting around the time of the faux-jump onset there was a greater increase in activity for the narrow goal than the wide goal. In contrast, the long head of the triceps (TLong, shoulder and elbow extensor) exhibited similar temporal structure and activity magnitudes for the narrow and wide goals (Figure 5B). Across the population of muscle samples for monkey M, 75% of samples had significantly greater activities for unperturbed reaches to the narrow goal than the wide goal in the movement epoch (two-sample t-test, p<0.01) with the average activity for the narrow goal being 15% greater than the activity for the wide goal (Figure 5C significant muscles are black filled circles). In contrast, there were no significant differences in activities for the narrow and wide goal in activity recorded from monkey C (Figure 5C, grey open circles).

**Figure 5.**
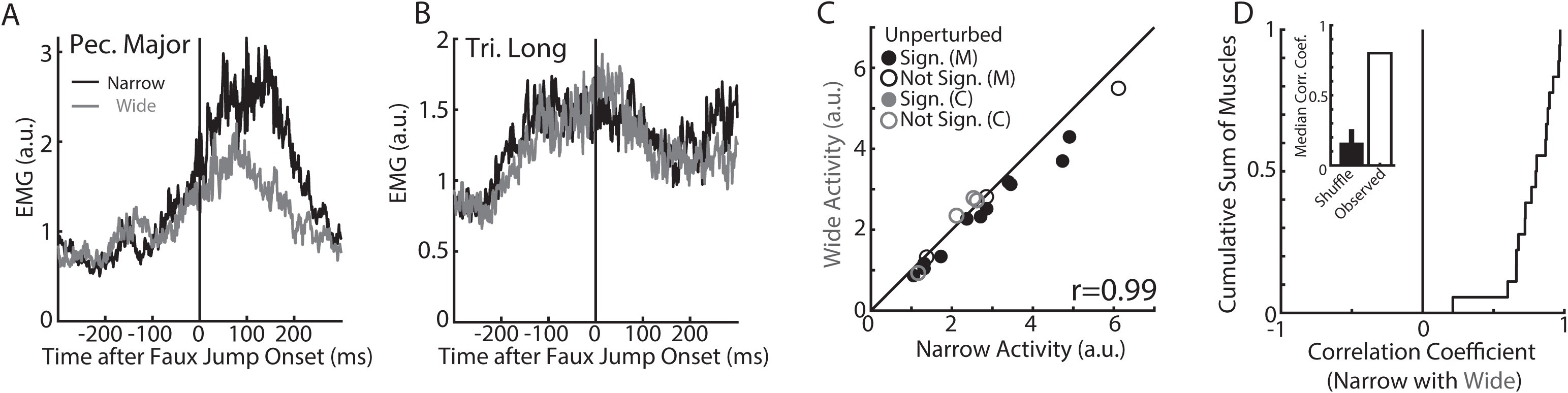
Muscle activity during unperturbed reaches to the narrow and wide goals for the cursor-variant of task. A) Average activity of the pectoralis major muscle for reaches to the narrow and wide goals. Activity are aligned to faux jump onset. B) Same as A) for the long head of the triceps. C) Comparison of the mean narrow and wide goal activities across muscles inside the 400ms epoch centered on the faux jump onset. ‘r’ denotes Pearson’s correlation coefficient. Monkey M and C are denoted by the black and grey markers. Filled circles denote muscles with significantly different activities for the narrow and wide goals. D) Cumulative sum of the temporal correlation coefficients between the narrow and wide goal activities across muscles (Monkey M and C pooled). Same 400ms epoch as C). Inset shows the median correlation coefficient from the shuffled distribution (mean ± standard deviation) and the observed median coefficient.

The magnitudes of the muscle activities were highly correlated between the narrow and wide goals (Pearson’s correlation coefficient r=0.99, all muscles included from monkey M and C). Furthermore, there was a strong temporal correlation between muscle activities for the narrow and wide goal reaches with a median correlation coefficient of 0.80 across muscles, which was significant (shuffle r=0.16, p<0.001; Figure 5D). Thus, muscle activity was largely similar between unperturbed reaches to the narrow and wide goals with a slight bias towards greater activity for reaches to the narrow goal for monkey M. Figures 6A and B show the change in activities (Δ EMG) for PM and TLong for cursor-jump trials (monkey M). For both muscles, activity started to differentiate between jump directions starting in <100ms. Activity also appeared to differentiate based on goal shape starting ∼100ms after the jump with greater change in activity for the narrow goal than the wide goal. We averaged across muscle samples and found the population activity increased from baseline for the narrow goal starting ∼89ms after the cursor jump (Figure 6C, bottom). However, we were unable to detect an onset for the wide goal. The population response differentiated based on the goal shape almost immediately with an onset detected at 89ms (Figure 6C, top). Similar results were found with monkey C except that the population signals started to differentiate from baseline (116ms, Figure 6D bottom) and between goal shapes (122ms, Figure 6D top) later which may be due to the noisier nature of surface EMG. Similar trends were found when analyzing onsets of individual muscles (Figure 6E).

**Figure 6.**
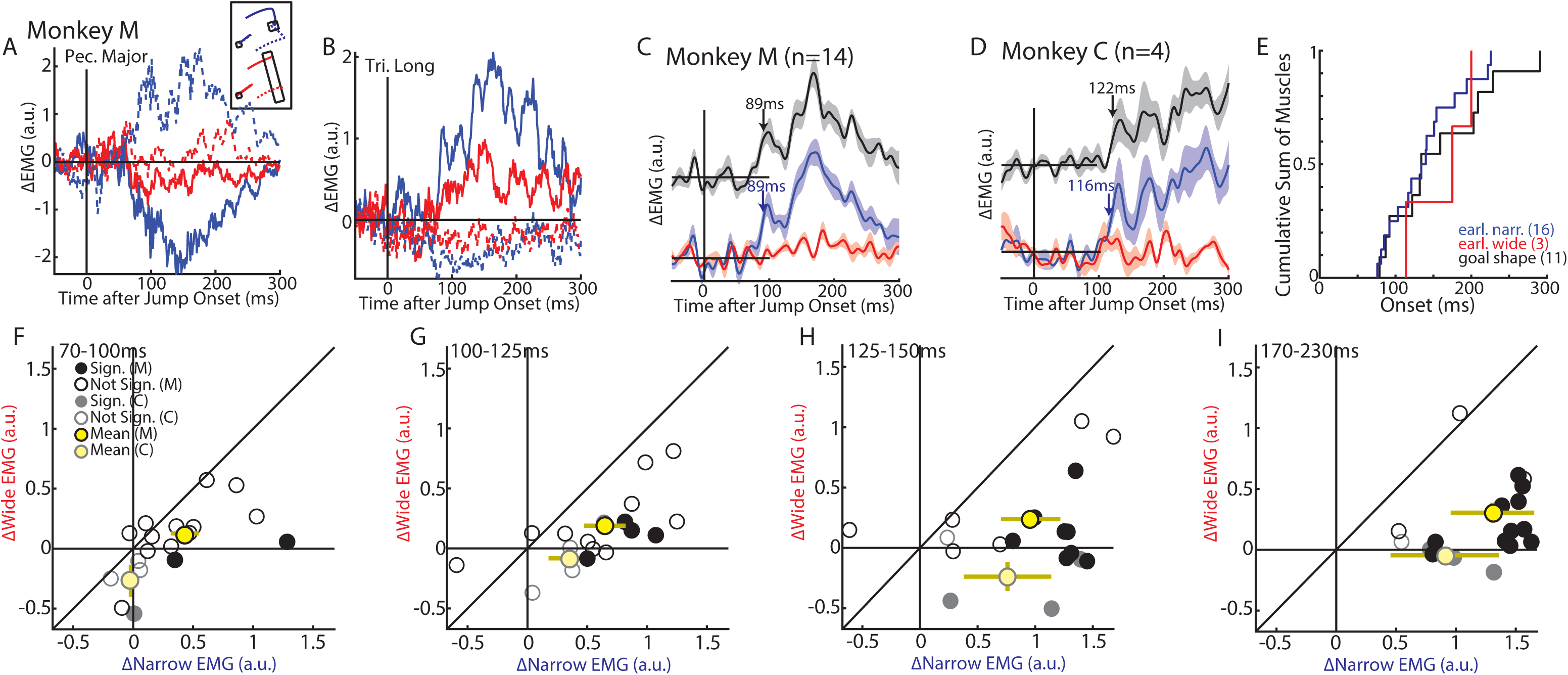
Muscle activity in response to the cursor jumps. A) The change in activity for the pectoralis major muscle in response to the cursor jumps when reaching for the narrow and wide goals. B) Same as A) for the long head of the triceps. C) Group average change in muscle activity to the cursor jumps for the narrow (blue, bottom) and wide goals (red, bottom). The resulting difference signal between the activities for the narrow and wide goals is shown in the black trace (top). Muscle activities were averaged across their preferred directions. Blue and black arrows denote when a significant increase in activity from baseline started for the narrow goal reaches and difference signal, respectively. No onset was detected for the wide goal reaches. D) Same as C) for Monkey C. E) Onsets for individual muscles presented as a cumulative sum. Numbers in brackets reflect the number of muscle samples with a detectable onset. Muscles recorded from Monkey M and C pooled. F) Comparison between the absolute change in muscle activities for the narrow and wide goals in the 70-100ms epoch. Muscles recorded from Monkey M and C are denoted in the black and grey markers. Yellow circles and bars denote the means and standard deviations for each monkey. Filled circles denote muscle samples that had significantly different activities for the narrow and wide goals. G-I) same as F) except for the 100-125ms (G), 125-150ms (H) and 170-230ms (I) epochs.

Next, we compared each muscle’s response in its preferred direction for the narrow and wide goals in the epochs of 70-100ms, 100-125ms, 125-150ms, and 170-230ms (Figure 6F-I). For each muscle we applied a two-way ANOVA with time (levels: four epochs) and goal shape (levels: narrow and wide) as factors and found 89% of samples had a significant interaction effect between time and goal shape (p<0.05, Bonferroni correction factor of 2). Post-hoc two-sample t-tests revealed, 17%, 27%, 61%, and 78% of samples had significantly different muscle responses in the 70-100ms, 100-125ms, 125-150ms and 170-230ms epochs, respectively (p<0.05, Bonferroni correction factor 4, Figure 6F-I filled circles). On average, the activity for the wide goal was 88%, 78%, 86% and 81% smaller than the activity for the narrow goal in the 70-100ms, 100-125ms, 125-150ms, and 170-230ms epochs, respectively. We examined group-level responses across trial-averaged muscle responses by applying a two-way repeated-measures ANOVA with time (levels: four epochs) and goal shape (levels: narrow and wide) as factors. We found a significant main effect of goal shape (F(1,17)=58.1 p<0.001) and an interaction between goal shape and time (F(3,51)=17.9, p<0.001). Post-hoc paired t-tests confirmed that responses for the narrow goal were significantly greater than for the wide goal in all epochs (70-100|100-125|125-150|170-230: df =11 (all epochs), t(11)=3.5| t(11)=8.3| t(11)=7.8| t(11)=9.7, p<0.01|p<0.001|p<0.001|p<0.001, Bonferroni correction factor 4). Collectively these results indicate muscle activity in response to visual feedback of the limb differentiates based on goal shape within ∼90-120ms.

## Experiment 2: Goal redundancy and feedback responses to mechanical loads

We modified the above reaching task to probe feedback responses to mechanical loads. We increased the size of the redundant dimension of the goal to better differentiate corrections to the narrow and wide goals (Figure 1B). We also adopted a shape for the wide goal that was similar to an arrowhead so that edges of the goal were in closer proximity to the monkey, thus making it easier to reach on perturbation trials. For monkeys M|A|T|C, we recorded 10|11|8|9 behavioural sessions of the animals performing the task on separate days. Again, monkeys were able to perform the task with high efficiency (success rates monkeys M|A|T|C: narrow targets=96|95|95|96%, wide targets=97|100|97|91%).

Figure 7A shows the hand paths for monkey M to the narrow (left) and wide (right) goals from one recording session. Similar to the previous task, there was more variability in the reaches to the wide goal (Figure 7B, C). Across sessions we found no significant different in endpoint position for monkey M and T, whereas there was a 0.7cm change in endpoint positions for monkeys A (t(10)=6.1 p<0.001) and C (t(8)=5.8, p<0.001). The variability of reach endpoints were 2.2|1.4|1.4|2.3 times greater for the wide goal than for the narrow goal for monkeys M|A|T|C, respectively (Figure 7E; paired t-test monkey M|A t(9)=5.3|t(10)=4.1|t(7)=5.2|t(8)=7.3, p<0.001|p=0.002|p=0.001|p<0.001).

**Figure 7.**
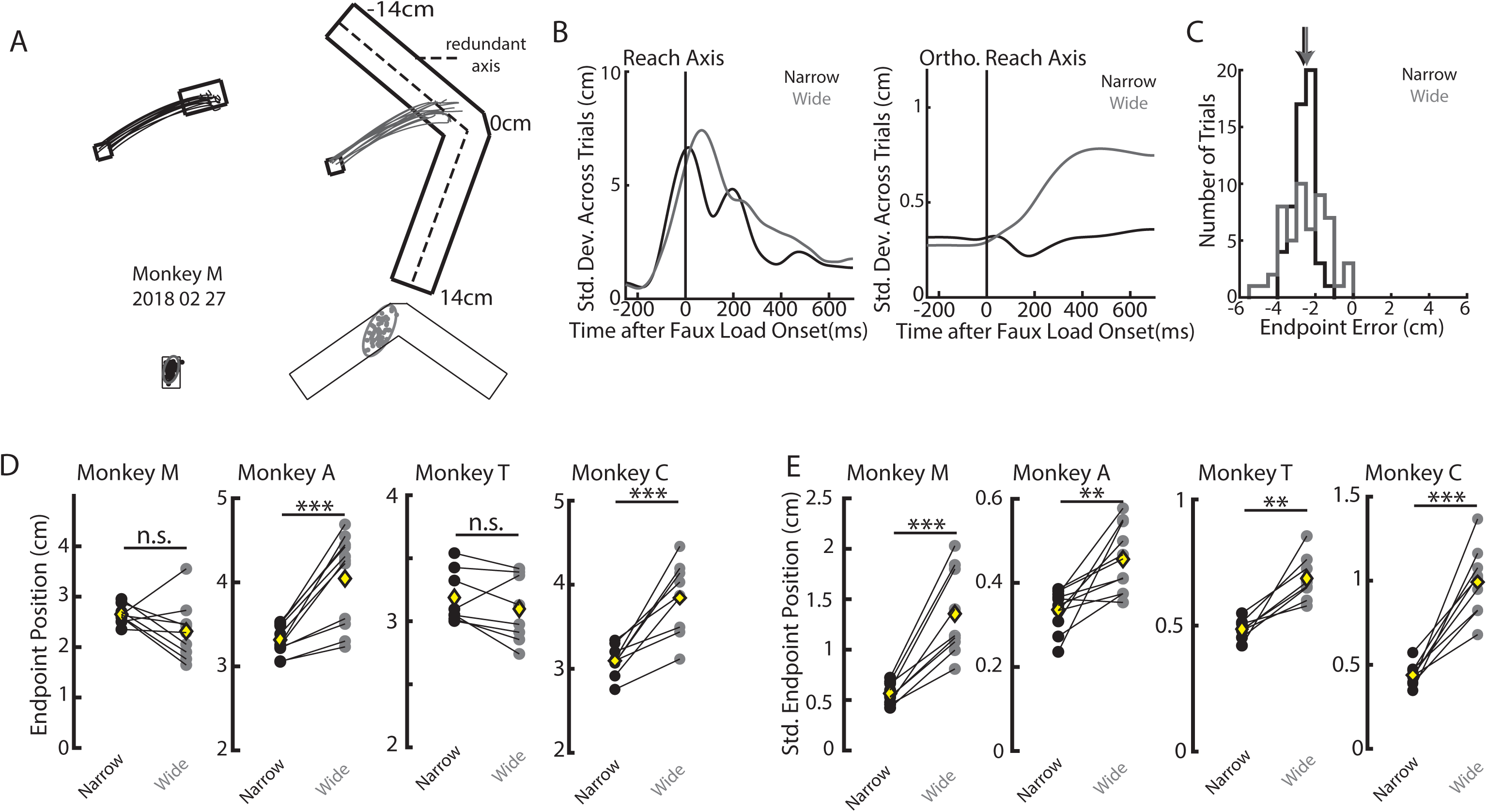
Example kinematics for unperturbed trials for the mechanical-load variant of task. A) Hand paths from one session for Monkey M to the narrow (left) and wide (right) goals. Dashed line on the wide goal denotes the redundant axis. B) Standard deviations of the hand position across trials for the narrow and wide goal reaches. C) Reach endpoint histograms for reaches to the narrow and wide goals from the same session as A). Zero denotes the middle of the redundant axis (A, middle of dashed line). Arrows denote the means of the distributions. D) The mean endpoint position for the narrow and wide goals across all recording sessions from both monkeys. Yellow diamonds denote the means across sessions. E) Same as D) for standard deviation of the endpoint position. ** p<0.01, ***p<0.001.

Figures 8A and B (inset) show the unperturbed hand velocities during the same recording session along the reach and orthogonal reach axes, respectively. In contrast to Experiment 1, there was a small decrease in the peak hand velocity in both directions for the wide goal as compared to the narrow goal. The hand speed was not significantly different for monkeys M (narrow 0.45m/s, wide 0.44m/s, paired t-test t(9)=1.5, p=0.2) and T (narrow 0.38m/s, wide 0.36m/s, t(7)=2.3 p=0.05), however there was a significant reduction for the wide goal for monkey A (narrow 0.52m/s, wide 0.47m/s, t(10)=8.0 p<0.001) and a significant increase for the wide goal for monkey C (narrow 0.36m/s, wide 0.38m/s, t(8)=2.6, p=0.03).

**Figure 8.**
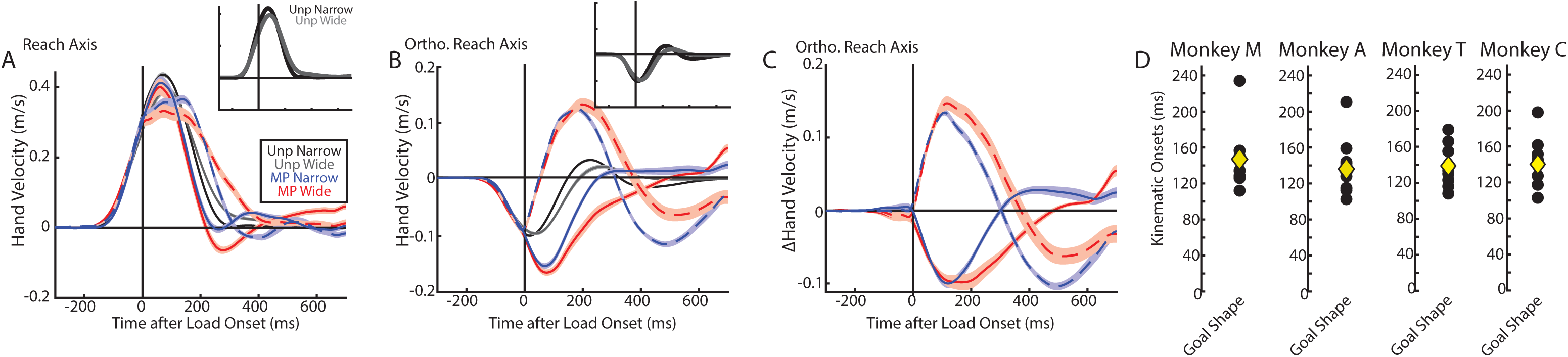
Example hand velocity profiles for the mechanical-load variant of task. **A**) The hand velocity along the reach axis for the narrow and wide goals from the same recording session as Figure 7A). Velocity was aligned to the load onset. Inset shows the unperturbed reaches to the narrow and wide goals. Unp Narrow and Unp Wide: Unperturbed reaches to the narrow and wide goals, respectively. MP Narrow and MP Wide: Mechanical load trials for reaches to the narrow and wide goals, respectively. Solid and dashed lines denote mechanical loads away from the body and towards the body, respectively. B) Same as A) for the hand velocity along the orthogonal reach axis. C) The change in the hand velocity on mechanical-load trials for the narrow and wide goals. D) Time when corrections differentiated based on goal shape across sessions. Yellow diamonds are the means across sessions.

Next, we examined how goal redundancy affected corrective responses for the unexpected mechanical loads. Figure 9A shows monkey M’s hand paths on mechanical-load trials. There is a clear correction present when the monkey was reaching for the narrow goal, whereas when reaching for the wide goal monkeys exhibited greater variability and corrected less for the mechanical loads (Figure 9B, C). Across sessions, there was a greater change in reach endpoints (change relative to unperturbed trials) for the wide goal than the narrow goal (Figure 9D, F; paired t-tests: mechanical loads away from/towards body monkey M t(9)=7.8| t(9)=4.9, monkey A t(10)=9.5| t(10)=20, monkey T t(7)=3.6|t(7)=7.6, monkey C t(8)=6.1|t(8)=11.7, p<0.01 for all comparisons). There was also greater variability in the reach endpoints on perturbation trials in both directions (Figure 9E, G; monkey M t(9)=8.3|t(9)=3.0, p<0.001|p=0.01, monkey A t(10)=5.4|t(10)=4.5, p<0.001|p=0.001, monkey T t(7)=4.8|t(7)=5.4, p=0.002|p=0.001, monkey C t(8)=4.9| t(8)=4.4, p=0.001|p=0.002).

**Figure 9.**
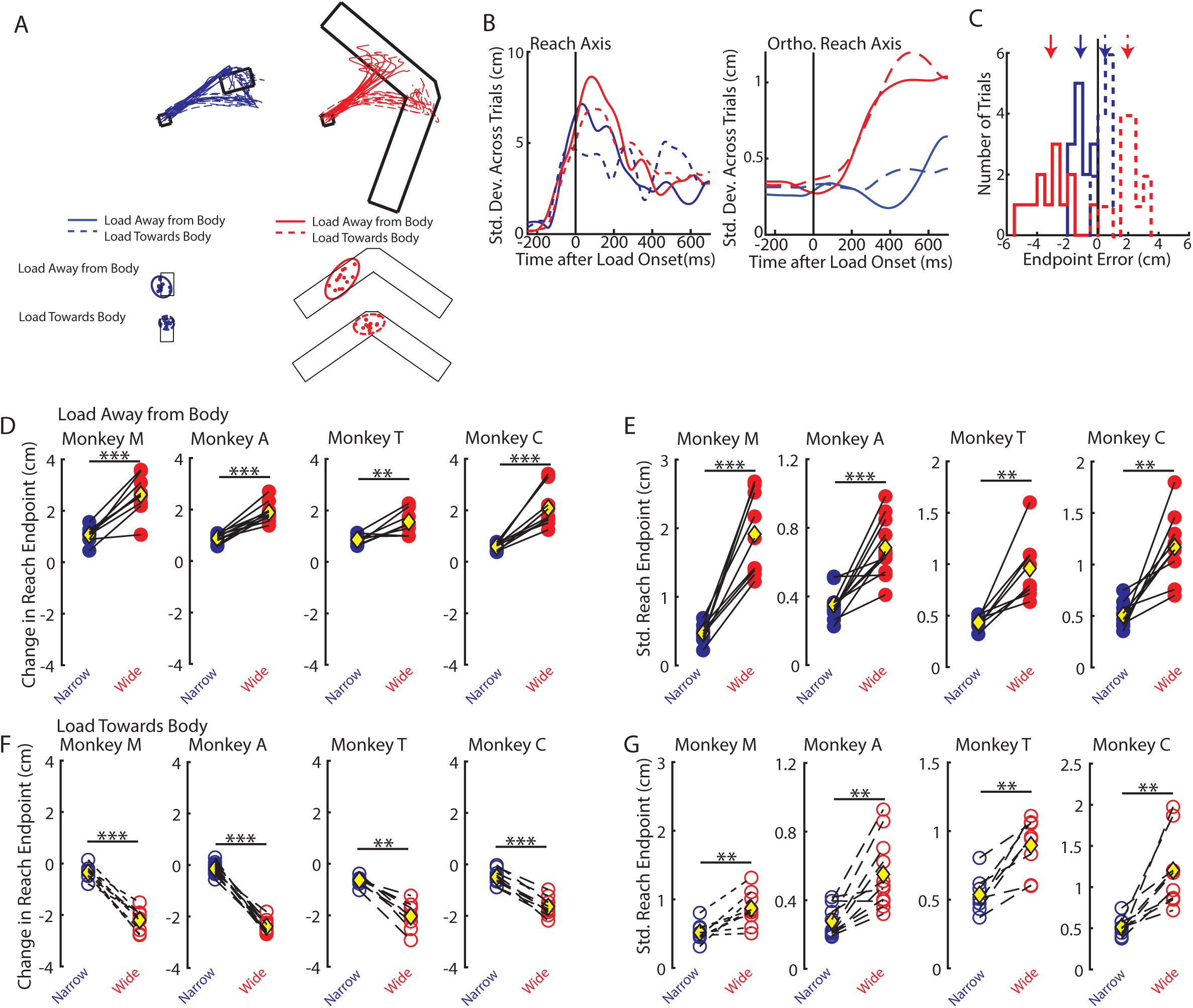
Example kinematics on mechanical-load variant of task. A) Top: Hand paths from Monkey M for mechanical-load trials. Same session as Figure 8A. Bottom: Reach endpoints and the 95% confidence ellipse. Solid and dashed lines denote mechanical load away from the body and towards the body, respectively. B) Standard deviations of the hand position across trials for the narrow and wide goal reaches. C) The change in reach endpoint histograms (change relative to unperturbed reach trials). Zero denotes the mean of the reach endpoints on unperturbed trials (Figure 8C arrows). Arrows denote the means of the distributions. D) The mean change in reach endpoint for loads away from the body across all recording sessions. E) Same as D) for the standard deviation in reach endpoints. F-G) Same as D-E) for loads towards the body. ** p<0.01, ***p<0.001.

For the mechanical-load trials, kinematic changes were primarily in the orthogonal reach axis though changes could also be detected in the reach axis (Figure 8A, B). Detecting the earliest kinematic correction to a mechanical load is difficult as the limb is already moving due to the momentum from the load (Figure 8C). However, we could detect differences in corrections based on goal shape that started at 147ms|136ms|139ms|140ms for monkeys M|A|T|C, respectively (Figure 8D).

For muscle activity recorded from monkey M (indwelling), we pooled recordings over the course of eight behavioural sessions as the differences in the corrective responses were comparatively weaker than for the cursor perturbations. All muscle samples from monkey M were included as they all showed perturbation-related activity. For monkey C, we again recorded using surface electrodes over the course of 7 recordings sessions and kept 6, 4, 3, and 6 recording sessions for TLat, Br, Bi, and TLong, respectively.

Figure 10A and B shows the average muscle activity on unperturbed trials for PM and TLong aligned to the faux-load onset (monkey M). Both muscles had similar temporal structure for the goal shapes, however, for PM there was greater activity for the narrow goal than the wide goal. Across muscle samples, all samples from monkey M had significantly greater activities for unperturbed reaches to the narrow goal than the wide goal in the movement epoch (two-sample t-test, p<0.01) with the average activity for the wide goal being 17% smaller than the activity for the narrow goal (Figure 10C significant muscles are filled black circles). In contrast, no muscle samples collected from monkey C were significantly different between the narrow and wide goals (grey circles). The muscle activity magnitudes were highly correlated between the two goal shapes for both monkeys (Pearson’s correlation coefficient across all muscles r=0.98) and there were was a strong temporal correlation between activities for the narrow and wide goals (Figure 10D, median r=0.891), and the distribution across muscles was shifted more to the right than the shuffled distribution (Inset, r=0.49, p<0.001).

**Figure 10.**
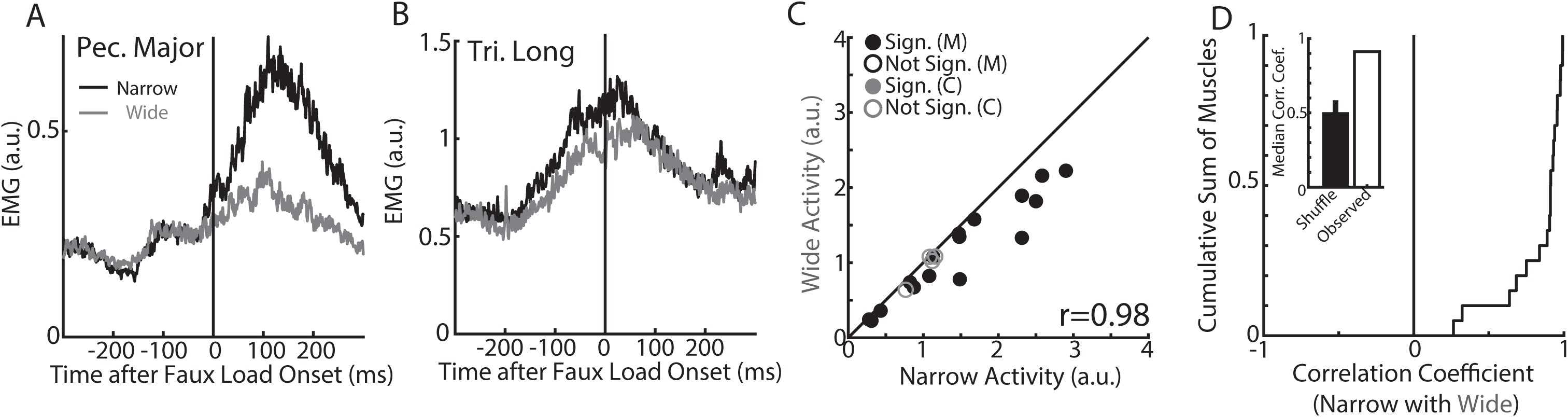
Muscle activity during unperturbed reaches to the narrow and wide goals for the mechanical-load variant of task. A) Average activity of the pectoralis major muscle for reaches to the narrow and wide goals. Activity are aligned to faux-load onset. B) Same as A) for the long head of the triceps. C) Comparison of the mean narrow and wide goal activities across muscles inside the 400ms epoch centered on the faux-load onset. ‘r’ denotes Pearson’s correlation coefficient. Monkey M and C are denoted by the black and grey markers. Filled circles denote muscles with significantly different activities for the narrow and wide goals. D) Cumulative sum of the temporal correlation coefficients between the narrow and wide goal activities across muscles (muscles from Monkey M and C pooled). Same 400ms epoch as C). Inset shows the median correlation coefficient from the shuffled distribution (mean ± standard deviation) and the observed median coefficient.

Changes in muscle activity in response to the mechanical loads started within ∼20-50ms of an applied load (Figure 11A-E). For monkey M, the population activity across load-sensitive muscles increased from baseline starting at 62ms and 64ms for the narrow and wide goals, respectively (Figure 11C). For monkey C, activity increased from baseline earlier at 33 and 26ms for the narrow and wide goals, respectively (Figure 11D). Activity differentiated based on goal shape starting at 78ms and 106ms for monkeys M and C, respectively (Figure 11C-E).

**Figure 11.**
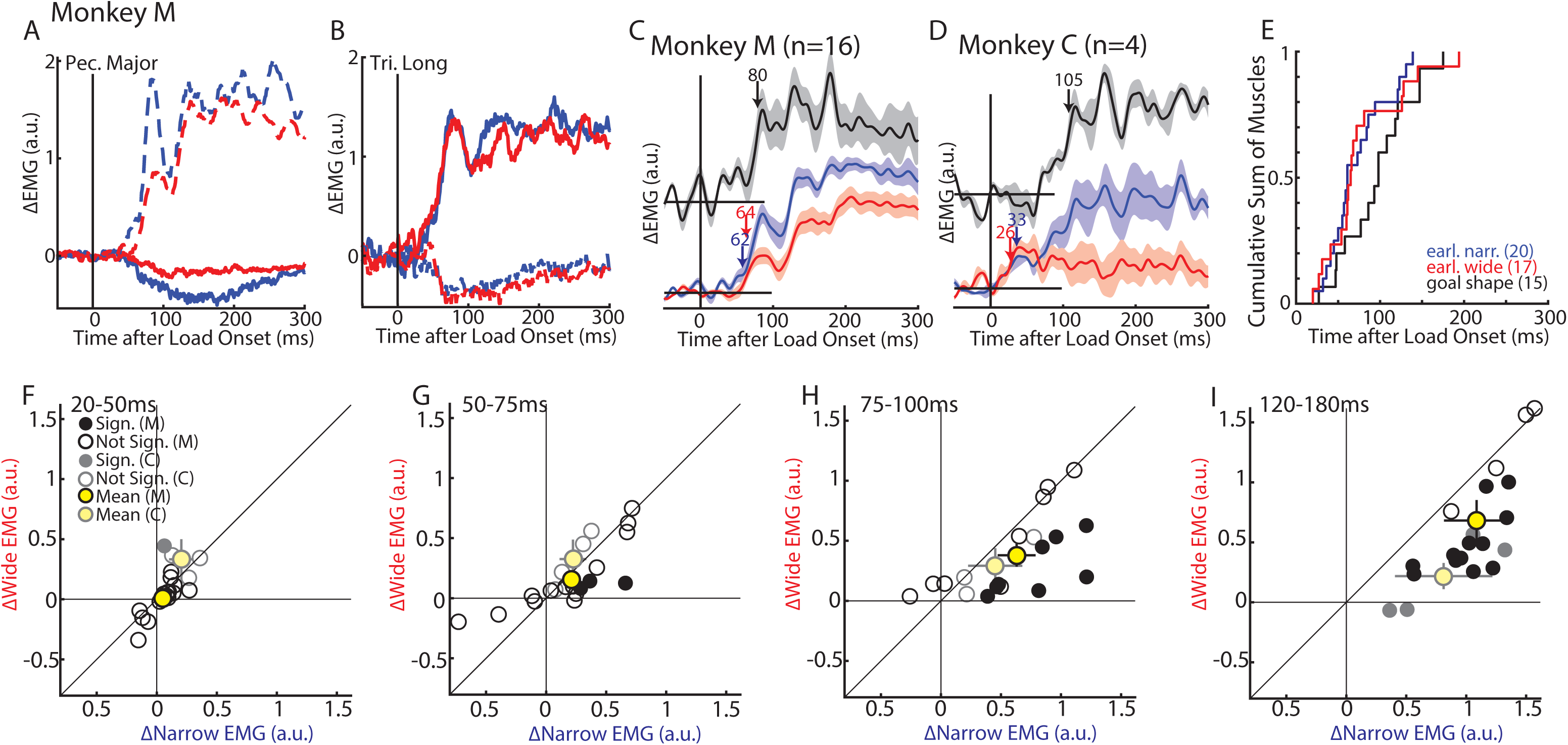
Muscle activity in response to the mechanical loads. A) The change in activity for the pectoralis major muscle in response to the mechanical loads when reaching for the narrow and wide goals. B) Same as A) for the long head of the triceps. C) Group average change in muscle activity for the mechanical loads for the narrow (blue, bottom) and wide goals (red, bottom). The resulting difference signal between the activities for the narrow and wide goals is shown in the black trace (top). Muscle activities were averaged across their preferred directions. Blue and red arrows denote when a significant increase in activity from baseline (500ms before mechanical load) started for the narrow and wide goal reaches, respectively. Black arrow denotes when a significant increase in activity from baseline started for the difference signal. D) Same as C) for Monkey C. E) Onsets for individual muscles presented as a cumulative sum. Numbers in brackets reflect the number of muscle samples with a detectable onset. Muscles from Monkey M and C were pooled. F) Comparison between the absolute change in muscle activities for the narrow and wide goals in the 20-50ms epoch. Muscles recorded from Monkey M and C are denoted in the black and grey markers. Yellow circles and bars denote the mean and standard deviation for each monkey. Filled circles denote muscle samples that had significantly different activities for the narrow and wide goals. G-I) same as F) except for the 50-75ms (G), 75-100ms (H) and 120-180ms (I) epochs.

Next, we compared each muscle’s response in the epochs of 20-50ms, 50-75ms, 75-100ms, and 120-180ms (Figure 11F-I). A two-way ANOVA with time and goal shape as factors found 85% of muscles samples had a significant interaction effect. Post-hoc two-sample t-tests revealed 6%, 17%, 47%, and 94% of samples had significantly different muscle responses in the 20-50ms, 50-75ms, 75-100ms and 120-180ms epochs, respectively (two sample t-tests p<0.05, Bonferroni correction factor 4, Figure 11F-I filled circles). On average, the activity for the wide goal was 13%, 12%, 39% and 43% smaller than the activity for the narrow goal in the 20-50ms, 50-75ms, 75-100ms and 120-180ms epochs, respectively. We applied a two-way RM ANOVA with time (levels: four epochs) and goal shape (levels: narrow and wide) to the trial-averaged muscle responses. We found a significant main effect of goal shape (F(1,19)=22.3 p<0.001) and an interaction between goal shape and time (F(3,57)=17.8, p<0.001). Post-hoc paired t-tests confirmed that responses for the narrow goal were significantly greater than for the wide goal in the 75-100ms (t(19)=3.2, p=0.004) and 120-180ms (t(19)=5.1, p<0.001) epochs, but not significantly different in the earlier epochs (20-50ms t(19)=0.3 p=0.7; 50-75ms t(19)=0.5 p=0.6).

## Discussion

Exploiting redundancies is an important feature of many motor control theories including OFC (Bernstein, 1967; Scholz et al., 2000; Todorov and Jordan, 2002; Latash, 2012). Several studies demonstrate how humans are capable of exploiting redundancies during motor actions (Todorov and Jordan, 2002; Diedrichsen, 2007; Mutha and Sainburg, 2009; Knill et al., 2011; Dimitriou et al., 2012; Nashed et al., 2012, 2014; Cluff and Scott, 2015). Here, we demonstrate that monkeys are able to exploit the spatial redundancy of a goal as previously observed in humans. Specifically, monkeys exhibited greater variability in their reach endpoints when reaching to a more spatially redundant goal. Rapid corrective responses to limb perturbations were also greatly attenuated to exploit goal redundancy with observable changes in muscle activity starting in <100ms.

Although it is impossible to know for certain whether the monkey recognized that they could reach anywhere on the wide targets, there are several lines of evidence that indicate monkeys exploited goal redundancy similar to humans and OFC models. First, OFC predicts that trial-by-trial variability even on unperturbed trials should be larger for goals with greater spatial redundancy with variability constrained along the redundant axis of the goal (Knill et al., 2011; Nashed et al., 2012). Previous studies in humans (Knill et al., 2011; Nashed et al., 2012; Cross et al., 2019) and the current study in monkeys demonstrate similar OFC-like structure with variability growing throughout the duration of the reach and culminating in greater variability in the reach endpoints along the redundant axis of the goal.

Second, OFC models predict that corrections to external perturbations should be smaller for more spatially redundant goals provided the perturbation is along the redundant axis. Previous studies demonstrate that humans share this feature in their corrective responses to visual and mechanical perturbations (Knill et al., 2011; Nashed et al., 2012; Cross et al., 2019). Here, we demonstrate monkeys also show smaller corrective responses when reaching for the wide goal. This was evident as a 2-4cm shift in the reach endpoints on perturbation trials from where monkeys were reaching on unperturbed trials. In contrast, there was <1cm shift in endpoints when monkeys were reaching for the narrow goal on perturbation trials. Endpoints on perturbation trials were also more variable when reaching for the wide goal indicating monkeys did not simply learn to reach for a particular goal location on perturbation trials. Collectively, these results argue that that monkeys in the present study understood goal redundancy with behaviour comparable to human performance and did not simply learn an arbitrary mapping between sensory stimuli and behavioural response that mimicked the expected behaviour.

Monkeys also generated muscle activity patterns to the mechanical loads that were similar to the OFC model and humans. OFC models predict an initial increase in control output in response to the mechanical load regardless of goal shape reflecting that the controller must counteract the external load to stabilize the limb (Nashed et al., 2012). Control output differentiates based on goal shape later with greater activity for the narrow goal to generate the necessary kinematic correction. Muscle activity evoked by the mechanical loads exhibit similar patterns as the control output with an initial increase starting at ∼50ms followed by differentiation based on goal shape starting at ∼70ms in both humans (Nashed et al., 2012) and monkeys (present study). Other studies have also found humans and monkeys exhibit similar timing for when corrective responses to mechanical loads are modulated by different contexts including for limb physics (Kurtzer et al., 2008; Pruszynski et al., 2011), task instruction (Hammond, 1956; Evarts and Tanji, 1976; Pruszynski et al., 2008, 2014; Omrani et al., 2014), and adaptation (Cluff and Scott, 2013; Maeda et al., 2018, 2020) which all start 60-70ms after the onset of the load. This similarity highlights how monkeys are a useful model to investigate the neural circuits that underlie flexible feedback processing during motor actions.

Our results contrast with previous findings by Bizzi and colleagues ( 1982, 1984) who found monkeys correct back to the original trajectory when encountering an assistive mechanical load. One possible reason is that monkeys may have learned an implicit timing constraint for when the arm should arrive at the goal and thus resisted the applied load to prevent arriving too early. Indeed, humans also show similar corrections but only when given a timing constraint (Cluff and Scott, 2015).

One limitation in our findings was that muscle activity was greater in amplitude for unperturbed reaches to the narrow goal than the wide goal, particularly for monkey M. Presumably, this increased activity was also present on mechanical-load trials and poses a potential problem for interpreting onsets to mechanical loads due to a gain-scaling effect where muscle activity evoked by a load scales with the size of the background muscle activity (Marsden et al., 1976; Bedingham and Tatton, 1984; Matthews, 1986; Stein et al., 1995; Pruszynski et al., 2009). Thus, greater muscle activity for the narrow goal following a mechanical load could simply reflect a gain-scaling effect. However, we believe this is unlikely as the effect of gain scaling typically only influences muscle activity within 20-50ms after an applied load (Pruszynski et al., 2009; Nashed et al., 2012) whereas in our study activity differentiated based on goal shape at ∼70ms. Second, we did not detect a significant difference in muscle activity for monkey C on unperturbed trials using surface electrodes and still found similar muscle timing.

Studies highlight that primary motor cortex (M1) is involved with generating flexible muscle responses to sensory feedback. M1 receives rich proprioceptive feedback with responses that start within ∼20ms of an applied load (Conrad et al., 1974, 1975; Wolpaw, 1980; Fromm et al., 1984; Bauswein et al., 1991; Picard and Smith, 1992; Herter et al., 2009; Takei et al., 2018; Heming et al., 2019; Cross et al., 2020, 2021). Importantly, proprioceptive feedback responses in M1 are modulated by several behavioural factors including limb physics (Pruszynski et al., 2011), prior instruction (Evarts and Tanji, 1976; Pruszynski et al., 2014), and task engagement (Omrani et al., 2014) within 50ms of an applied load. The flexible responses in M1 preceded the corresponding change in muscle activity by ∼10ms consistent with the conduction delay between M1 and the periphery (Cheney and Fetz, 1984; Lemon et al., 1986). Thus, if M1 is involved with generating proprioceptive feedback responses that exploits goal redundancy than M1 activity reflecting goal redundancy should emerge ∼60ms after the load onset.

However, it is likely that other brain areas also contribute to generating muscle responses that exploit goal redundancy. Premotor cortex, somatosensory cortex, parietal area 5and the cerebellum all project to M1 (Jones et al., 1978; Porter and Lemon, 1993; Dea et al., 2016) and rapidly respond to proprioceptive feedback within <70ms with activity patterns that are context dependent (Wolpaw, 1980; Lamarre et al., 1983; Strick, 1983; Chapman et al., 1984; Pruszynski et al., 2011, 2014; London and Miller, 2012; Omrani et al., 2014, 2016). However, context is not homogenously shared across these areas as premotor cortex, M1 and cerebellum exhibit activity patterns consistent with implementing a control policy whereas somatosensory and parietal areas exhibit patterns of activity consistent with state estimation (Strick, 1983; Omrani et al., 2016). These results are also supported by a recent study examining temporary inactivation of a subset of these areas (Takei et al., 2021). Thus, given that goal redundancy is a property of the control policy, we predict context-dependent responses will emerge first in premotor cortex, M1 and the cerebellum.

Monkeys also generated muscle activity patterns to the cursor jumps that were similar to humans and OFC models. Unlike mechanical loads, cursor jumps do not require the controller to counteract an external load as the disturbance is only a kinematic error. Thus, control output of the OFC model should be unaffected by the cursor jump when reaching for the wide goal. Similarly, muscle activity of humans (Cross et al., 2019) and monkeys are largely unchanged by a cursor jump when reaching to a wide goal resulting in activity that differentiates based on goal redundancy ∼90ms after a cursor jump. This differentiation is also unlikely due to a gain-scaling effect as background muscle activity does not appear to affect correction strength for visual perturbations (Franklin et al., 2017).

It is less clear how visual feedback is processed by fronto-parietal circuits and how behavioural context influences visual processing in these areas. A common assumption is that visual feedback is processed by posterior parietal cortex which is then sent towards frontal circuits including M1 and premotor cortex (Goodale and Milner, 1992; Desmurget et al., 1999; Pisella et al., 2000; Gaveau et al., 2014). Thus, consistent with OFC visual feedback is processed initially by circuits involved with state estimation followed by circuits involved with implementing the control policy. However, there is evidence that visual feedback responses arrive first in premotor cortex (50-70ms) followed by M1 (70-100) and finally parietal area 5 (Cisek and Kalaska, 2005; Archambault et al., 2011; Ames et al., 2014; Stavisky et al., 2017; Cross et al., 2021). Thus, premotor cortex may generate the earliest muscle response to visual feedback rather than M1.

Alternatively the superior colliculus may be involved with generating rapid motor responses to visual feedback (Alstermark et al., 1987; Day and Brown, 2001; Pruszynski et al., 2010; Corneil and Munoz, 2014; Day, 2014; Cross et al., 2019; Kozak et al., 2019). Activity in the superior colliculus correlates with muscle activity of the upper arm during reaching (Werner, 1993; Werner et al., 1997; Stuphorn et al., 1999) and stimulation of the superior colliculus can evoke reaching-like behaviour (Philipp and Hoffmann, 2014). Further investigations are needed to elucidate the underlying neural circuits involved with generating rapid visual responses for which our behavioural task could be invaluable.

## Acknowledgements

We thank Kim Moore, Simone Appaqaq, Catherine Crandell, Jordan Miller, Ethan Heming, and Helen Bretzke for their laboratory and technical assistance. This work was supported by grants from the Canadian Institute of Health Research to SHS. KPC was supported by an Ontario Graduate Scholarship. SHS was supported by a GSK chair in Neuroscience.

